# Novel Bruton’s Tyrosine Kinase (BTK) substrates for time-resolved luminescence assays

**DOI:** 10.1101/2022.04.05.487161

**Authors:** Naomi E. Widstrom, Minervo Perez, Erica D. Pratt, Jason L. Heier, John F. Blankenhorn, Lindsay Breidenbach, Hannah Peterson, Laurie L. Parker

**Author notes:** **Corresponding Author** Laurie L. Parker,. These authors contributed equally (Naomi E. Widstrom, Minervo Perez). National Institutes of Health, National Cancer Institute, Chemical Biology Laboratory, 1050 Boyles St., Bldg. 538, Rm. 244, Frederick, MD 21702. Boston University, Department of Biomedical Engineering, 44 Cummington Mall, Boston, MA 02215.

## Abstract

Bruton’s tyrosine kinase (BTK) is a well-documented target for cancer therapeutics due to its role in B-cell signaling pathways. However, inhibitor design is hindered by lack of tools to assess kinase activity. We used in vitro phosphoproteomics to determine BTK’s substrate preferences and applied this information to our updated data processing pipeline, KINATEST-ID 2.1.0. This pipeline generates a position-specific scoring matrix for BTK and a list of candidate synthetic substrates, each given a score. Characterization of selected synthetic substrates demonstrated a correlation between KINATEST-ID 2.1.0 score and biochemical performance in *in vitro* kinase assays. Additionally, by incorporating a known terbium-chelation motif, we adapted synthetic substrates for use in an antibody-free time-resolved terbium luminescence assay. This assay has applications in high-throughput inhibitor screening.

## INTRODUCTION

Nonreceptor Bruton’s tyrosine kinase (BTK) sits at a node of immune signaling pathways, controlling both adaptive and innate immune cell function, proliferation and other key signaling pathways.^*1*^ In B-cells, BTK activity is required for maturation and B-cell receptor signaling, as well as mediating signals from chemokine receptors.^*2-4*^ In myeloid cells, BTK has a role in toll-like receptor signaling and modulating the innate immune response.^*5*^ Dysregulation of BTK is linked to a number of different B-cell leukemia and lymphomas.^*6*^ Additionally, BTK expression in tumor-associated macrophages within the tumor microenvironment is linked to promotion of cancer cell migration and metastasis.^*7*^ Due to these roles, BTK has been an attractive target for cancer therapeutics. To date, there are three FDA-approved BTK inhibitors for treatment of various B-cell malignancies: ibrutinib, acalabrutinib, and zanubrutinib.^*8*^ All three are irreversible inhibitors that covalently bind to cysteine 481 in the kinase domain of BTK, effectively inhibiting BTK activity. Although these inhibitors show promising results,^*9-12*^ disease relapse due to resistance remains an issue.^*13, 14*^ Therefore, there is a need to continue BTK inhibitor development, which relies on sensitive, high-throughput screens to determine kinase activity. Kinase activity can be assessed by a variety of means, such as radiometric or fluorescent based approaches, but these methods are higher cost or reliant on antibodies, making them less ideal for screening.^*15*^

Lanthanide luminescence assays are an option for kinase activity detection that have the advantage of being antibody free. Lanthanides such as terbium absorb light poorly on their own, but in the presence of a sensitizing chromophore, such as phosphotyrosine, terbium exhibits high luminescence with an increase in luminescent lifetime.^*16*^ To take advantage of these properties, multiple approaches have been described using a peptide incorporating a terbium-chelation motif and appropriate sensitizer to detect phosphorylation.^*17-20*^ Previously, we have adapted a sensitive time-resolved terbium luminescence assay into a format that works well for high-throughput drug screening.^*21-23*^ In this assay, a synthetic peptide substrate with a central tyrosine residue is phosphorylated by the kinase of interest. When phosphorylated, the peptide can better chelate the Tb^3+^ ion allowing tyrosine to sensitizes the Tb^3+^ luminescence. Tb^3+^ chelation and sensitization greatly enhances the luminescence intensity, which is especially pronounced when using time-resolved luminescence measurements. This allows for sensitive differentiation of the phosphorylated form of the synthetic substrate, providing a fast readout of kinase activity through straightforward liquid addition steps (without requiring complex handling). The peptide substrate used in this assay must contain a terbium-chelating motif, while also maintaining the required substrate motif to be phosphorylated by the kinase of interest. Therefore, we developed an *in silico* pipeline called KINATEST-ID to assist in designing such substrates by identifying kinase preferences and providing information on which positions are amenable to substitution of amino acids that facilitate chelation.^*22*^ However, design of these synthetic substrates was hindered by the lack of known BTK substrates. Here, we describe our work to determine the substrate profile of BTK using in vitro phosphoproteomics, updates to the tools used to analyze the substrate profile and design synthetic substrates (KINATEST-ID 2.1.0), and the optimization of novel synthetic BTK substrates for use in the time-resolved luminescence assay.

## RESULTS and DISCUSSION

### Design of BTK synthetic substrates

The KINATEST-ID approach uses information about a kinase’s known substrates to identify novel optimal substrate candidate sequences.^*22*^ However, very few substrates for BTK have been reported, and we required a larger knowledgebase in order to design novel substrates effectively. Therefore, we employed a modified phosphoproteomics workflow designed to determine the substrate profile of a kinase (similar to our previous reports for developing FLT3 substrates).^*24-26*^ In brief, we treated trypsin-digested cell lysate with phosphatase to remove endogenous modifications, followed by reaction with recombinant BTK for two hours. Phosphopeptides resulting from this reaction were enriched and identified using an Orbitrap Fusion mass spectrometer (Figure 1). The identified substrates were input into our updated data processing workflow, KINATEST-ID 2.1.0 (Figure 2). KINATEST-ID 2.1.0 is an updated R package of our previous KINATEST-ID workflow, comprising a full set of data handling and analysis steps. The updated v2.1.0 uses a Fisher Odds-based positional scoring matrix to calculate the odds ratio and significance of each amino acid at each position relative to the central phosphorylation site, as opposed to the standard deviation model we used previously.^*22*^ It also includes amino acids from -7 to +7 flanking the phosphorylation site, in order to provide information about whether resides on either end (−7 to -5 and +5 to +7) can be substituted easily with acidic residues to improve chelation of terbium for the time-resolved luminescence assay (however, it only uses positions -4 to +4 for predicted substrate scoring, since that region is typically employed in studies of tyrosine kinase substrate recognition using peptides).^*27*^ From the phosphoproteomics data, we found a total 68 nonredundant substrates in common between the three technical replicates after subtraction of substrates found in control samples. Representing the most robust observations, these substrates were used to calculate the position-specific odds ratio of each amino acid and generate a heatmap that illustrates the substrate motif identified through this *in vitro* phosphorylation experiment (Figure 3A).

**Figure 1.**
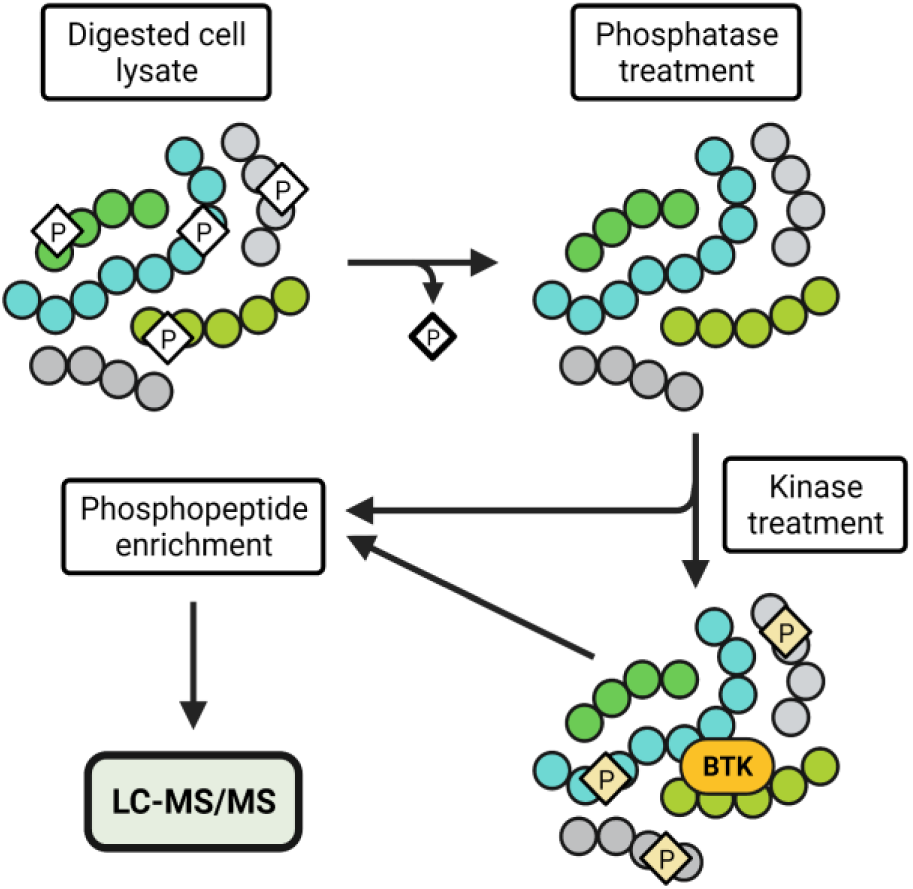
Schematic of phosphoproteomic workflow. KG-1 cells were lysed, trypsin-digested, and treated with alkaline phosphatase to create a natural peptide library. The peptide library was treated with recombinant BTK and phosphoenriched prior to LC-MS/MS analysis. Created with BioRender.com.

**Figure 2.**
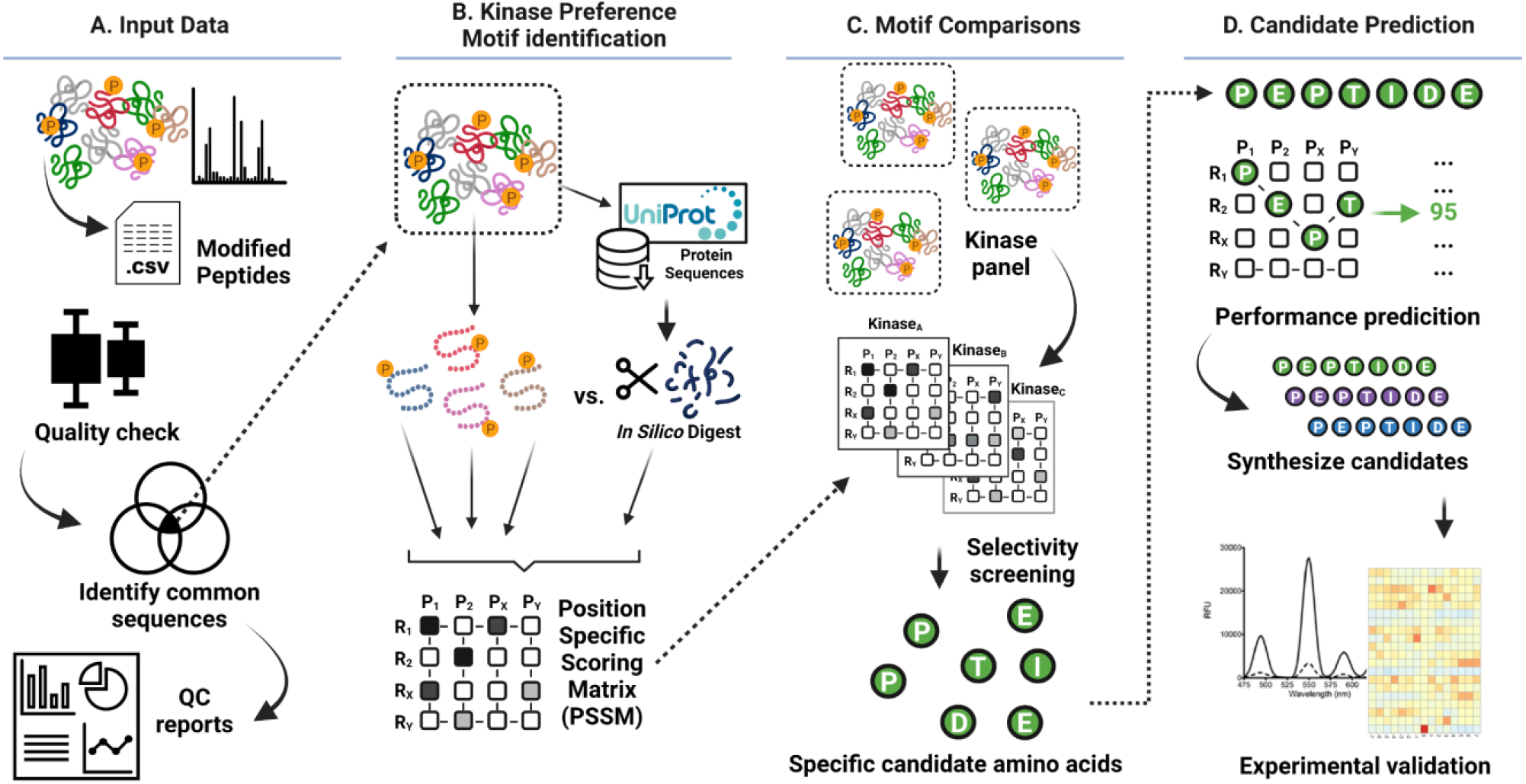
Schematic of KINATEST-ID 2.1.0. A.) Results lists of the phosphopeptides identified in the in vitro phosphoproteomics experiment are filtered to select the sequences robustly observed in common between replicates, as well as generate quality control reports. B.) Robustly observed sequences are used in combination with a surrogate “background” consisting of an *in silico* tryptic digest of protein sequences drawn from the FASTA file using UniProt identifiers linked to phosphopeptides observed, to generate a position-specific scoring matrix for the kinase of interest. C.) The target kinase preferences are screened against a panel of off-target kinases’ preferences to determine motif elements most likely to provide a selective substrate, identifying candidate amino acids and refining selectivity-focused motifs. D.) A table of sequences representing all permutations of candidate amino acids for the -4 to +4 positions is generated, each with a score for predicted biochemical performance. From here, candidate sequences are selected for synthesis and experimental characterization. Created with BioRender.com.

**Figure 3.**
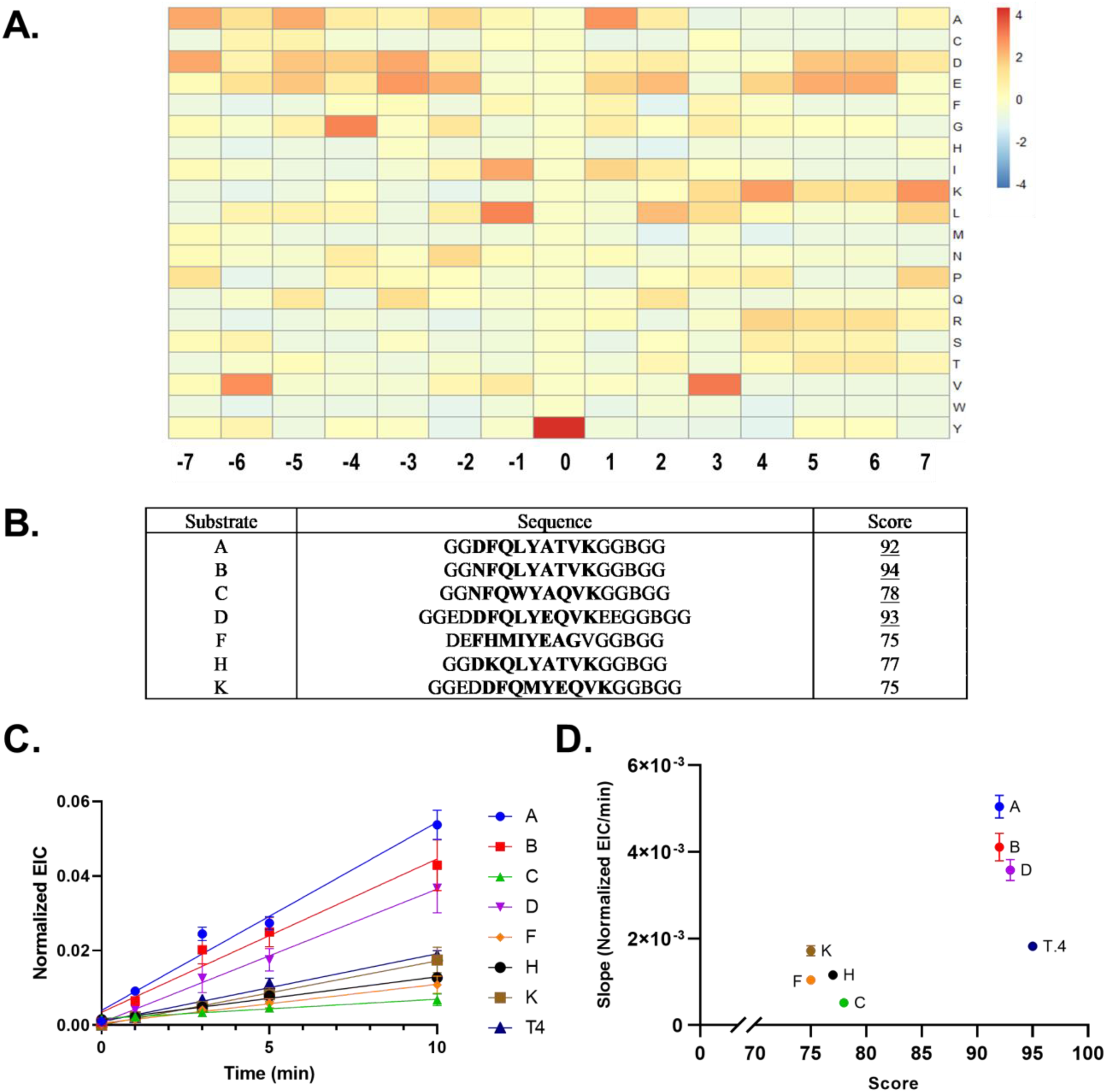
BTK phosphoproteomics results and synthetic substrate performance. A.) Heatmap showing amino acid frequency across positions relative to central tyrosine. B.) Synthetic substrate sequences and scores. B denotes biotinylated lysine; bold text denotes designed substrate portion containing the phosphorylation site. Underlined text denotes predicted active substrates. C.) Kinase assay with LC-MS-based readout of the 8 synthetic substrates in the presence of recombinant BTK. Product EIC (P) was normalized with substrate EIC (S) with Eq. 3. Data points are average of 3 replicates ± SD. D.) Comparison of KINATEST-ID score for each substrate vs. the slope of normalized EIC. Data points are average of 3 replicates ± SEM. Pearson correlation coefficient r = 0.7421with a p-value of 0.0350. EIC, extracted ion chromatogram.

Using the KINATEST-ID 2.1.0 *Generator* function, we generated candidate sequences according to BTK’s identified substrate motif. We selected seven candidate sequences to synthesize. The motif comparison *Screener* module (Figure 2C) was not considered in the design of these peptides, due to the goal of developing an *in vitro* assay in which only a single kinase would be present. Four sequences represented -4 to +4 from the central phosphorylation site (the core recognition motif^*27*^), and three were extended to -7 (and in one case also to +7) to incorporate chelating residues. Sequences were chosen to have a range of KINATEST-ID 2.1.0 scores (calculated just for the -4 to +4 segment, as mentioned previously, using a scoring function derived from the Fisher Odd’s table, Eq. 2) to test if there was a correlation between the sequence’s score and how well the substrate performed in a BTK kinase assay. The candidate BTK synthetic substrates were given letter names (A, B, C, D, F, H, K) and each synthesized with glycines flanking the substrate portion, and a biotinylated lysine near the C-terminus, to allow affinity capture of the peptides, followed by additional glycines. An initial *in vitro* kinase assay was performed with recombinant BTK, collecting and quenching four aliquots over 60-min incubation, with phosphorylation detected using an anti-phosphotyrosine antibody in an ELISA-based assay (Supplementary Figure S1). As a follow-up to that initial progress curve experiment, we used LC-MS to more thoroughly monitor linear product formation from each substrate from 0 to 10 mins (Figure 3C). We compared the KINATEST-ID 2.1.0 scores with the initial slope of phosphorylation using a Pearson correlation analysis and found that there is a statistically significant correlation between the score and the initial rate of phosphorylation of the substrates (Figure 4D). This suggests that when a relevant *in vitro* phosphoproteomics dataset is used to determine the Fisher’s Odds table and scoring function, the scores generated by KINATEST-ID 2.1.0 are useful to approximately predict the efficiency of a substrate. While all synthetic substrates would still need to be validated using a kinase assay, this information is useful in deciding what range of scores are most likely to indicate an efficient synthetic substrate. These findings demonstrate that we are able to successfully design synthetic substrates for BTK that perform well in *in vitro* BTK kinase assays.

**Figure 4.**
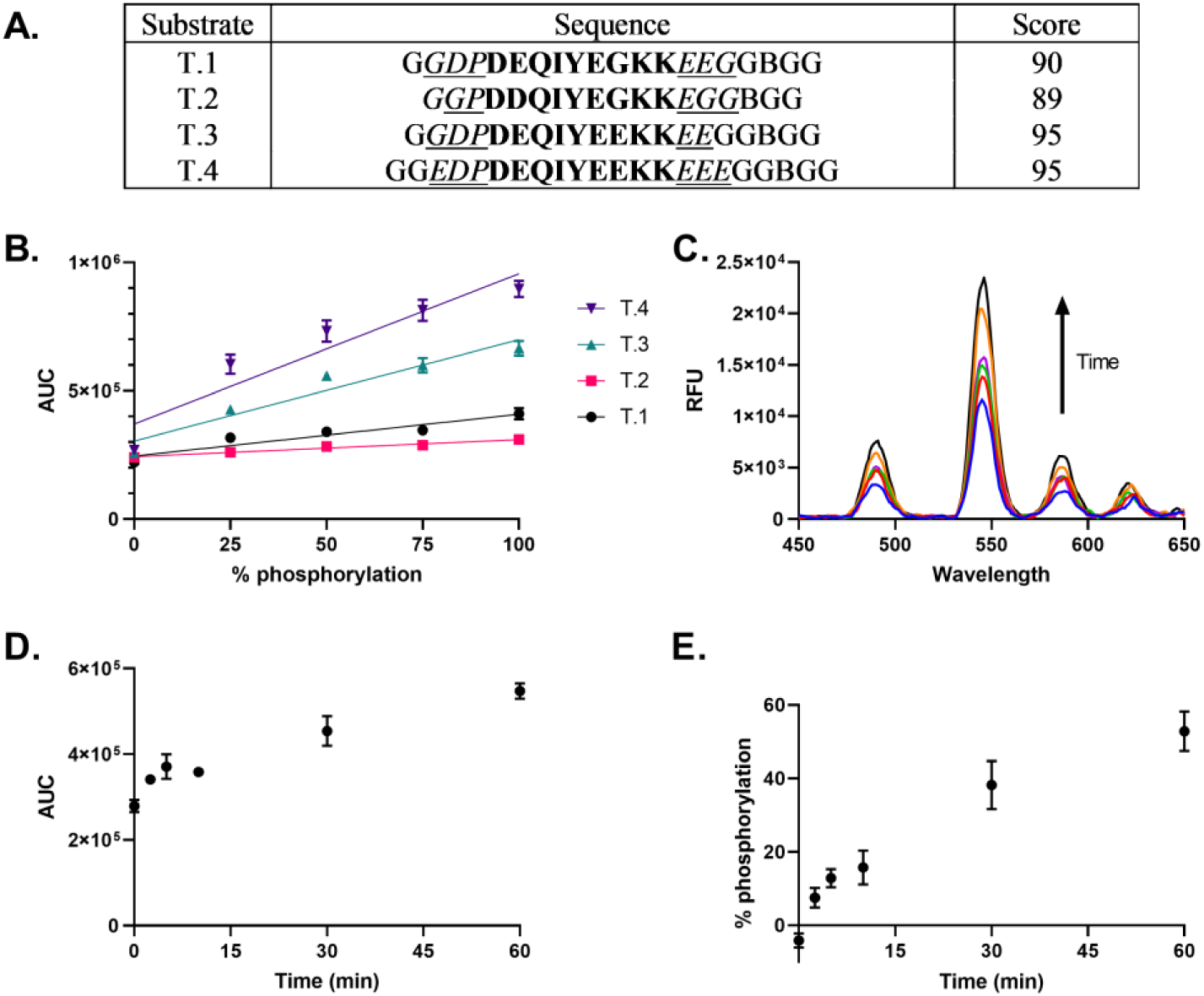
Characterization of terbium-chelating BTK synthetic substrates. A.) Sequences of terbium-chelating substrates and their KINATEST-ID V2.1.0 scores. B denotes biotinylated lysine; Bold text denotes designed substrate portion containing the phosphorylation site; Italic text denotes the -7 to +7 positions evaluated for compatibility of acidic residues with BTK’s substrate preferences. B.) Dynamic range of four synthetic substrates. Each peptide mixed with different ratios of its phosphorylated form to test the relative range and sensitivity. Data are shown as an average of six technical replicates ± SEM. C-E.) Terbium luminescence spectra (C) from *in vitro* BTK kinase assay with BTK synthetic substrate T.4 and area under curve (AUC) for each spectra (D). Percent phosphorylation of BTK synthetic substrate T.4 across time interpolated from regression line from standard curve (E). Kinase assay data points represent 6 replicates ± SEM. RFU, relative fluorescent units; AUC, area under curve.

### Design of terbium-chelating substrates

While several of the synthetic substrates showed promising results in the ELISA-based assay, the ELISA itself is not ideal for inhibitor screening, as its multiple wash and incubation steps make high-throughput testing difficult. Although sequence D was capable of chelating and sensitizing Tb^3+^ when phosphorylated, its signal to background for unphosphorylated vs. phosphorylated was too low to achieve useful dynamic range in the assay. Therefore, we focused attention on specifically designing BTK synthetic substrates for use in the sensitive, high-throughput terbium luminescence assay. Previously, we described the characterization of Syk Artificial Substrate peptide (SAStide), which facilitates terbium chelation when the tyrosine is phosphorylated, leading to luminescence enhancement.^*21*^ SAStide’s terbium binding motif was designed based on a the alpha synuclein sequence surrounding position Y125 (DPDNEAYEMPSEEG), which sensitizes terbium when phosphorylated.^*18*^ We noted that BTK prefers acidic amino acids at a number of positions, making it an ideal candidate to incorporate into the terbium motif. We chose one sequence frame based on the alpha synuclein sequence^*18*^, with a sequence predicted by KINATEST ID to be a BTK substrate (DEQIYEGKK) as the central -4 to +4 substrate portion, and altered the total number of acidic residues by substituting in glutamic acid residues for glycine to investigate the role of acidic amino acids in this motif.

These synthetic substrates were designated ‘T’ with variants 1 through 4: synthetic substrates T.1, T.2, T.3, and T.4 (Figure 4A). We tested the signal to background ratio by synthesizing a phosphorylated version of each substrate and mixing the two versions together in different ratios to create a standard curve from 0 to 100% phosphorylation (Figure 4B). These experiments were done in the same kinase assay reaction buffers as the actual reactions, to mimic conditions and background of a standard kinase assay. We found that while the background level of each substrate remained similar, by increasing the number of acidic residues we were able to increase the signal to background ratio for detecting phosphorylation and achieve a wide dynamic range. As BTK synthetic substrate T.4 was found to have the best dynamic range of the terbium-chelating synthetic substrates, we next tested its efficiency as a BTK substrate. We performed a kinase assay by adding 20 µM synthetic substrate T.4 to a kinase reaction mixture containing 20 nM recombinant BTK and quenching aliquots in urea at specific time points. Following addition of TbCl_3_, the time-resolved spectra were immediately measured (Figure 4C). Area under the curve (AUC) was calculated from the emission spectra (Figure 4D) and a standard curve containing 0 to 100% phosphorylated T.4 (Figure 4D) and used to interpolate the percent phosphorylation (Figure 4E). A clear progression of increased phosphorylation was observed over time and by 60 mins, approximately 50% of the synthetic substrate was phosphorylated, demonstrating that T.4 acts as an efficient substrate for BTK. In the subsequent experiment comparing initial rates for the non-Tb-sensitizing peptides and T.4 (Fig. 3C), its phosphorylation was not as efficient as some of the best substrates—however, T.4 was the preferred terbium-binding peptide with the highest signal to noise ratio (S/N) of the four peptides tested (3.34 for T.4, which was 2.5-fold higher than 1.29 for T.2). We felt this trade-off in substrate efficiency in order to achieve a viable lanthanide luminescence assay was acceptable. Overall, these findings demonstrate that the BTK synthetic substrate T.4 can be effectively used with the time-resolved luminescence assay to rapidly determine the activity of BTK.

In summary, we used phosphoproteomics to determine the substrate preference of BTK and optimized tools to use this information to design efficient synthetic substrates for BTK. We found a correlation between our KINATEST-ID 2.1.0 score and substrate performance, establishing that our process can approximate which synthetic substrates are likely to be efficient substrates of BTK, aiding in sequence selection for substrate design. BTK synthetic substrate A proved to be an effective substrate with both the ELISA and LC-MS readout. For designing terbium-chelating synthetic substrates for use in our time-resolved luminescence assay, we demonstrated that increasing the number of acidic residues outside the motif of amino acids preferred by BTK resulted in an increase in the dynamic range for time-resolved lanthanide luminescence detection of phosphorylation, without affecting the background signal. BTK synthetic substrate T.4 proved to be an efficient BTK substrate as well. This novel substrate provides an antibody-free assay that is rapid, sensitive, and requires only simple liquid addition steps, qualities ideal for the high-throughput screening necessary for BTK inhibitor testing. Additionally, the process described here of substrate discovery and synthetic substrate design can be easily applied to other tyrosine kinases to provide useful tools for kinase assay development.

## Supporting information

BTK KINATEST-ID output files

## ASSOCIATED CONTENT

### Supporting Information

The Supporting Information is available free of charge at: Materials and Methods, and supplementary figures (PDF)

KINATEST-ID 2.1.0 output files (ZIP file)

Mass spectrometry files (raw data, PEAKS Studio X pro output files, PEAKS ModExtractor script files) are deposited in the MassIVE repository (https://massive.ucsd.edu, accession #MSV000088429).

## AUTHOR INFORMATION

### Author Contributions

NEW, MP and LLP conceived the project. NEW, MP, JLH, HP and LB performed experiments. NEW, MP, JFB and EDP analyzed data. JFB designed and wrote initial R scripts, EDP wrote the updated and consolidated KINATEST-IDv2.1.0 R package. NEW, EDP and LLP wrote the manuscript.

### Notes

The authors declare no competing financial interests.

## ACKNOWLEDGMENT

This work was supported by the National Institutes of Health/National Cancer Institute through R01CA183571 and R33CA217780 (LLP), R01CA183571-S1 (MP), T32CA009138 (NEW and EDP), and T32AG029796 (JLH and JFB). The table of contents graphic was created with BioRender.com.

---

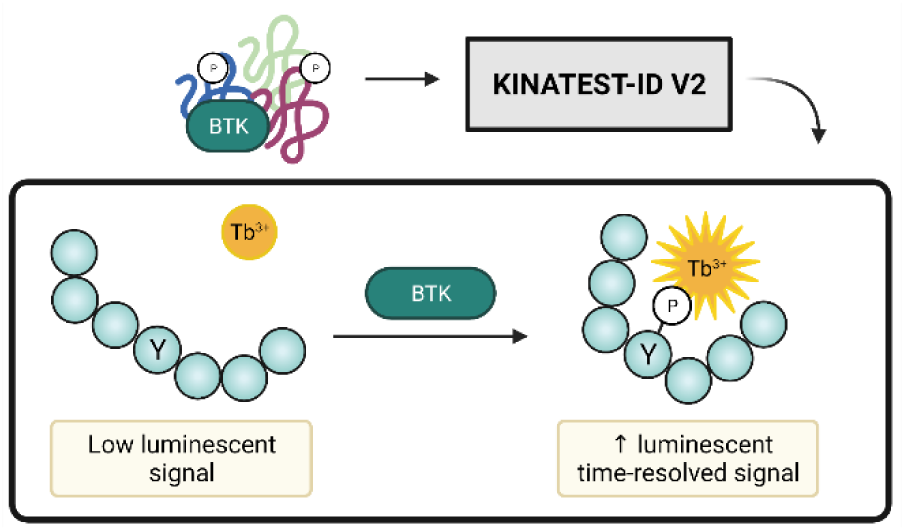

