## Supplemental Materials for "Novel Bruton’s Tyrosine Kinase (BTK) substrates for time-resolved luminescence assays"

‡These authors contributed equally.

**MATERIALS and METHODS**

**Cell culture and natural peptide library preparation.** KG-1 cells (ATCC) were maintained in IMDM media (Gibco) supplemented with 20% heat inactivated fetal bovine serum (FBS), 1% penicillin/streptomycin in 5% CO_2_ at 37°C. Endogenous peptide sample preparation was carried out as described previously.*^1^* In brief, cells were lysed, alkylated, and trypsin digested prior to being treated with alkaline phosphatase to remove endogenous phosphate groups. For three replicate samples, 1.5 µg recombinant BTK (SignalChem) was added to half of the endogenous peptide sample in a kinase reaction mixture (50 mM Tris HCL, pH 7.5, 10 mM MgCl_2_, 1 mM DTT, 1 mM Na_3_VO_4_ and 2 mM ATP) and incubated for two hours at 37°C. The other half of the sample was processed and analyzed as a control for effectiveness of the phosphatase reaction. All samples were phosphoenriched using PolyMac kits (Tymora Analytical) prior to mass spectrometry analysis.

**LC-MS/MS data acquisition.** LC-MS/MS data acquisition was carried out as described previously with a modification to the LC elution time interval.*^1^* In brief, samples were reconstituted in LC/MS solution (98/2/0.5%; H₂O/ACN/formic acid (FA)) and loaded on a ThermoScientific Easy NanoLC LC 1000 system. The mobile phase consisted of 0.1% formic acid in ultra-pure water (Solvent A) and 0.1% formic acid in acetonitrile (Solvent B). Samples were run over a linear gradient (5-30% solvent B; 80 minutes) with a flow rate of 200 nL/min into a high resolution Orbitrap Fusion Tribrid Mass Spectrometer, using the same parameters as previously described.*^1^*

**Data processing.** PEAKS Studio X pro (Bioinformatics Solutions Inc) was used to process and identify the sequences of the modified peptides. Following sequence identification, we exported the protein-peptides list for input into the PEAKS ModExtractor script*^2^* (according to its instructions, available in the repository housing the script <https://gitlab.com/jackbrennan07/peaks-modextractor>) which combines all input files into a single output file containing the identified peptides, UniProtIDs, modification site and A score. This file was used as the input for the R package KINATEST-ID 2.1.0.

Building on earlier work*^1, 3^*, KINATEST-ID 2.1.0 (<https://github.com/llparkerumn/KINATESTIDv2>) integrates curated collections of known endogenous substrate sequences and data from phosphoproteomics-informed kinase assays to develop an *in silico* screening library. The position-specific amino acid preferences contained in the library inform the design of high-performance artificial peptide substrates. Using KINATEST-ID 2.1.0, the initial peptide list (n = 5647) was filtered for sequences containing a phosphotyrosine (n = 314). For increased robustness and specificity, the peptide list was additionally filtered for sequences present in all three kinase-treated technical replicates but absent in negative controls (n = 68).

These sixty-eight input peptides were used to construct a position-specific scoring matrix (PSSM). The frequency of each amino acid was calculated for flanking residues -4 to +4 from the central tyrosine. Frequencies were converted to odds ratios applying the Fisher's Exact Test and used to generate the PSSM. A peptide *P* can be described by the PSSM as follows:

$$P_{PSSM}= \left[ \begin{matrix} a_{11} & a_{12} & \ldots& a_{1j} \\ a_{21} & a_{22} & \ldots& a_{2j} \\ \vdots& \vdots& \ldots& \vdots\\ a_{i1} & a_{i2} & \ldots& a_{ij} \end{matrix} \right]$$

(1)

Where *a_ij_* represents the Fisher odds ratio for amino acid *i* at flanking position *j*. The background frequencies for calculating the odds ratios were based on the input peptides. The full protein sequences were downloaded for each UniprotKB/Swiss-Prot ID found in the 68 sequence input dataset (https://www.uniprot.org/help/uniprotkb). *In silico* tryptic digestion was used to generate a control list of tyrosine-containing polypeptides (n = 3301) which should have been present in the original endogenous peptide library but were not modified by the kinase of interest. Candidate amino acids with a_ij_ > 1 were selected at each flank position based on: 1) p ≤ 0.05 or odds ratio > 2) the number of kinases in the screening library which *disfavored* that residue (odds ratio < 1).

Candidate amino acids were then permuted to generate a list of potential kinase-specific artificial peptide substrates. The total score *s* for each peptide *P* is the product of the odds ratio of each amino acid in the sequence:

$$s_{P}= \prod_{j = 1}^{J} a_{ij} where i = (A, C, \ldots, Y)$$

(2)

ROC analysis was also performed using the input peptides and control list to estimate a cutoff score for predicted peptide activity. Samples were bootstrapped using the R package 'cutpointr' (maximize metric) to calculate a more robust cutoff estimate. Generally, potential artificial peptide substrates scores above the cutoff are predicted to be biochemically active (i.e. phosphorylated) for that kinase. For interpretability, reported scores are log-transformed and scaled between 0 to 100.

**Peptide synthesis and purification.** Peptides were synthesized on Symphony X Peptide Synthesizer (Protein Technologies) on a 50 µmol scale with Rink amide resin (Protein Technologies). Fmoc-protected amino acids (Protein Technologies) were coupled in activating solution of HCTU in N-methylmorpholine (NMM) and dimethylformamide (DMF) in two 10-minute couplings. For biotinylated lysine the coupling time was 2 hours, and phosphotyrosine the coupling time was 8 hours. Fmoc group deprotected using 20% piperidine in DMF. Peptides were cleaved from resin using a cleavage cocktail of 94% trifluoroacetic acid (TFA), 2.5% H_2_O, 2.5% ethane dithiol and 1% Triisopropylsilane (TIS) followed by precipitation in ice cold diethyl ether and lyophilization. Peptides were then purified by reverse phase HPLC (Agilent 1200 Series Infinity LCMS) to >90% purity (Supplemental Figure 2). Synthetic substrates were dissolved in 100 mM (4-(2-hydroxyethyl)-1-piperazineethanesulfonic acid) (HEPES) pH 7.5 for LC-MS readout assays, and in phosphate buffered saline (PBS) for ELISA readouts and terbium time-resolved assays.

**Synthetic substrate *in vitro* assay with HPLC-MS readout.** Reactions (230 µL total volume) using recombinant BTK (SignalChem) were performed in triplicate at 25°C in kinase buffer (25 mM HEPES pH 7.5, 10 mM MgCl₂, 100 µM ATP, 3 mM DTT, 3 µM Na_3_VO_4_) with a BTK concentration of 10 nM. After incubating BTK in kinase buffer for 15 min., reactions were initiated with the addition of substrate (20 µM). Reaction aliquots (50 µL) were withdrawn and quenched with 10 µL 20% TFA (3.3% TFA in the quenched solution). Sample aliquots were run on HPLC (Agilent 1200 series) using a C18 Agilent Zorbax column (2.1 x 250mm, 5-micron) at 0.25 mL/min with acetonitrile increasing from 10% to 30% over 40 min. Reaction progress was analyzed via MS (Agilent 6130A) through integration of extracted ion chromatograms (EIC) corresponding to products (P) and substrates (S) (Agilent ChemStation) and normalized according to equation:

(3)

$Normalized EIC=\frac{product EIC}{(product EIC+substrate EIC)}$

Slope of the timepoints was calculated using GraphPad Prism, and a Pearson correlation analysis used to calculate the correlation between slope and KINATEST-ID 2.1.0 score.

**Synthetic substrate *in vitro* assay with ELISA readout.** Recombinant BTK (SignalChem) was diluted 10X in (20 mM MOPS pH 7.5, 1 mM EDTA, 0.01% Brij-35, 5% Glycerol, 0.1% beta-mercaptoethanol and 1 mg mL^-1^ bovine serum albumin (BSA)). Diluted BTK was incubated with reaction mixture (25 mM HEPES pH 7.5, 10 mM MgCl₂, 100 µM ATP, 3 mM DTT, 3 µM Na_3_VO_4_) to a final concentration of 20 nM for 15 mins at 37°C. Peptide substrate was added to a final concentration of 37.5 µM. Sample aliquots were quenched by combining 1:1 with 30 mM EDTA at timepoints. Chemiluminescent detection of phosphorylation was performed as previously described.*^1^*

**Synthetic substrate *in vitro* assay with terbium time-resolved readout.** Recombinant BTK (SignalChem) was added to a final concentration of 20 nM in kinase reaction mixture (25 mM HEPES pH 7.5, 10 mM MgCl_2_, 1 µM Na_3_VO_4_, 100 µM ATP, 0.05 µg µL^-1^ BSA) and incubated at room temperature for 15 mins. The reaction was started by adding the synthetic substrate at a final concentration of 20 µM to the reaction mixture. At timepoints 2.5, 5, 10, 30, and 60 mins a 20 µL aliquot was removed and quenched 1:1 in 6 M urea in a 384-well black plate. 10 µL of terbium luminescent mix (final concentration 100 mM NaCl, 200 µM TbCl_3_) was added to each well for a final well volume of 50 µL. The “zero” time point contained BTK storage buffer only (no enzyme). Time-resolved emission spectra were collected on Synergy Neo2 (Biotek) plate reader with monochromator excitation of 266 nm, time-resolved delay of 50 µsec and collection time of 1000 µs. Emission spectra collected between 450-650 nm with 1 nm step and a gain of 230. Area under the curve calculated using GraphPad Prism for each spectrum. Signal to noise ratio (S/N) was calculated by taking the average value of 100% phosphorylation AUC divided by the average value of 0% phosphorylation AUC for each substrate.

**Standard curves for terbium time-resolved assay.** Synthetic phosphorylated and unphosphorylated peptide were added in ratios of 0, 25, 50, 75, and 100% phosphorylation with a final concentration of 20 µM peptide. Procedure and buffer solutions used were the same as for the kinase assay, minus the addition of the kinase itself.

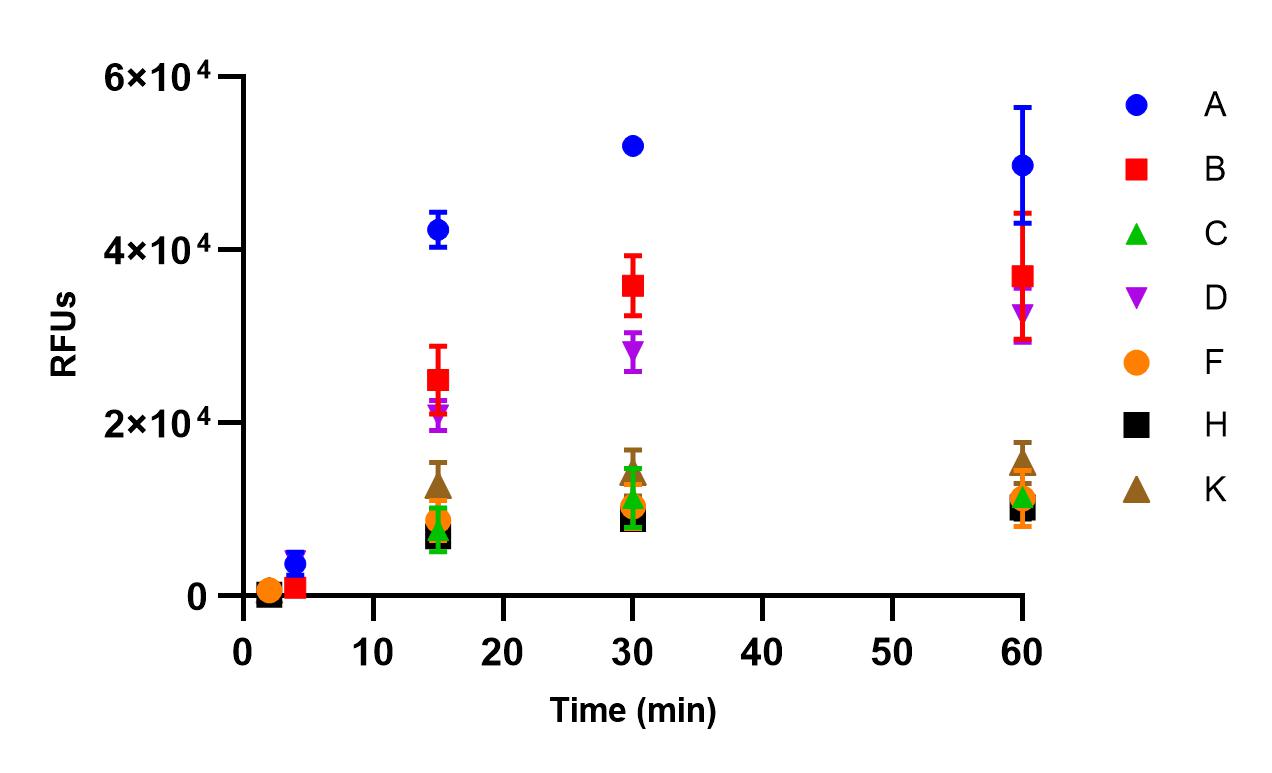

**Figure S1.** **Kinase assay with ELISA-based readout**. Progress curves for phosphorylation of BTK synthetic substrates A, B, C, D, F, H, and K (37.5 µM) in the presence of recombinant BTK. Aliquots of the reaction mixture were quenched 1:1 in EDTA at timepoints 4, 15, 30, 60 mins (A, B, C, D) or 2, 15, 30, 60 mins (F, H, K). Data points from A, B, C, D average of 3 replicates, data from F, H, K average of 6 replicates ± SEM. RFU, relative fluorescent units.

**Figure S2. Characterization of Peptides**

**A.** BTK synthetic substrate A (GGDFQLYATVKGGK_BIOTIN_GG)

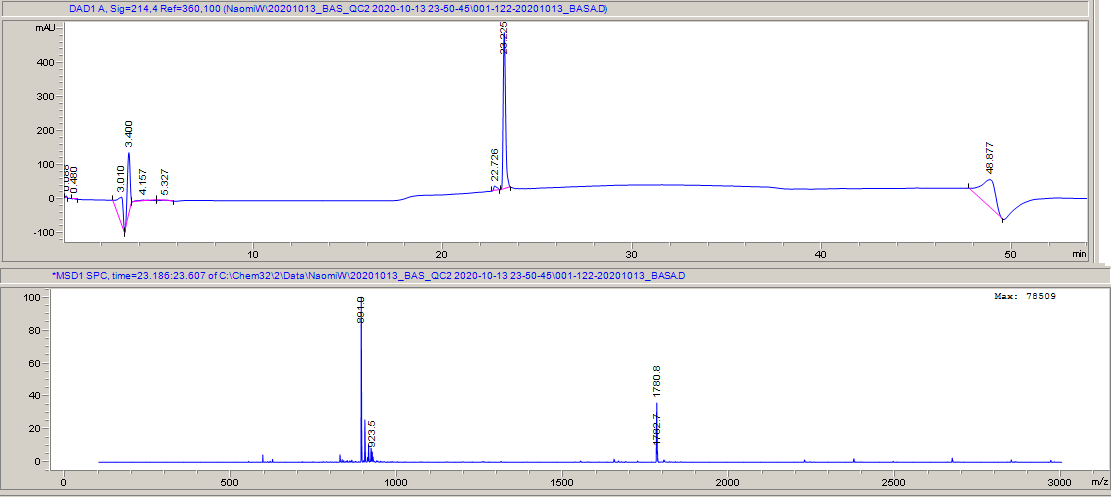

**B.** BTK synthetic substrate B (GGNFQLYATVKGGK_BIOTIN_GG)

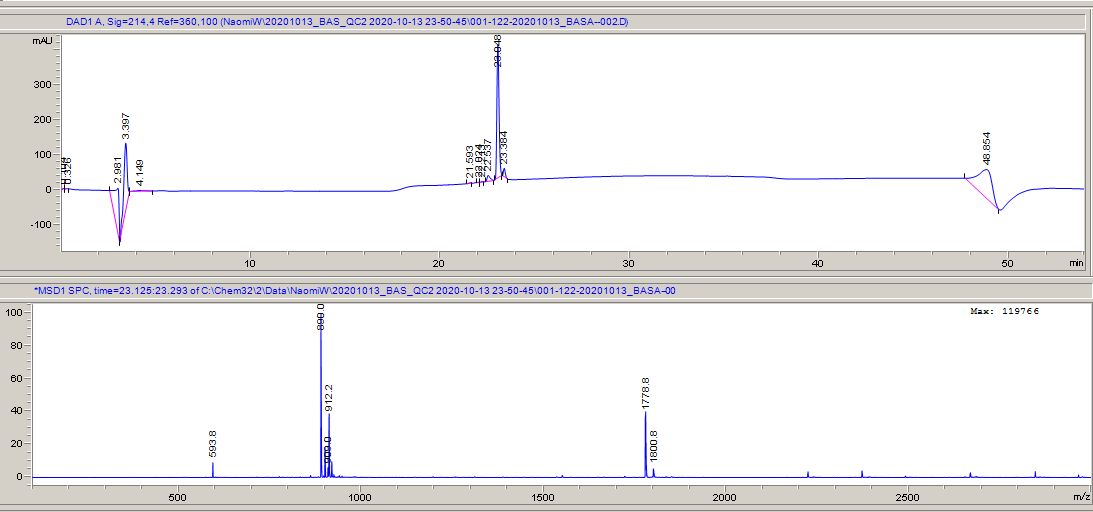

**C.** BTK synthetic substrate C (GGNFQWYAQVKGGK_BIOTIN_GG)

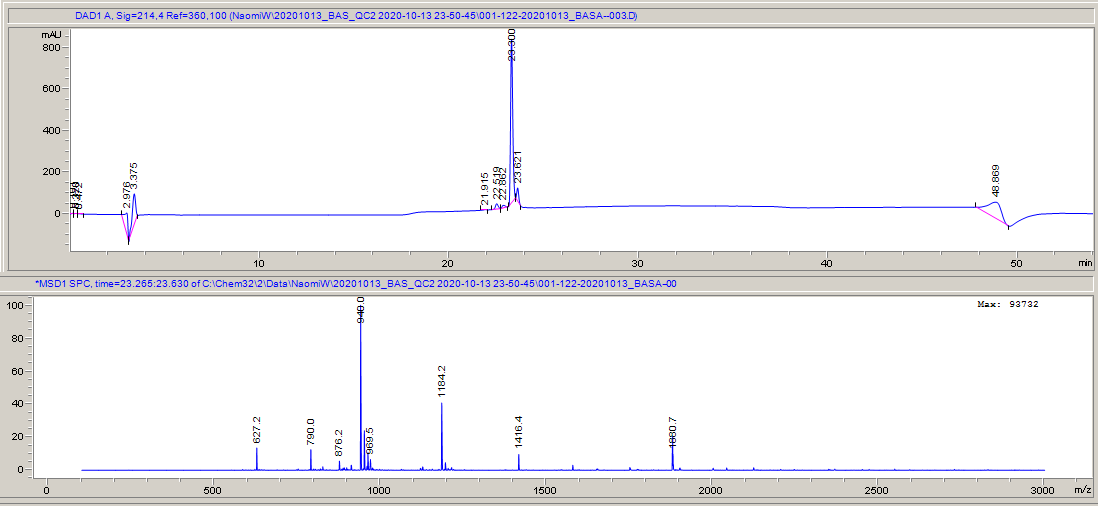

**D.** BTK synthetic substrate D (GGEDDFQLYEQVKEEGGK_BIOTIN_GG)

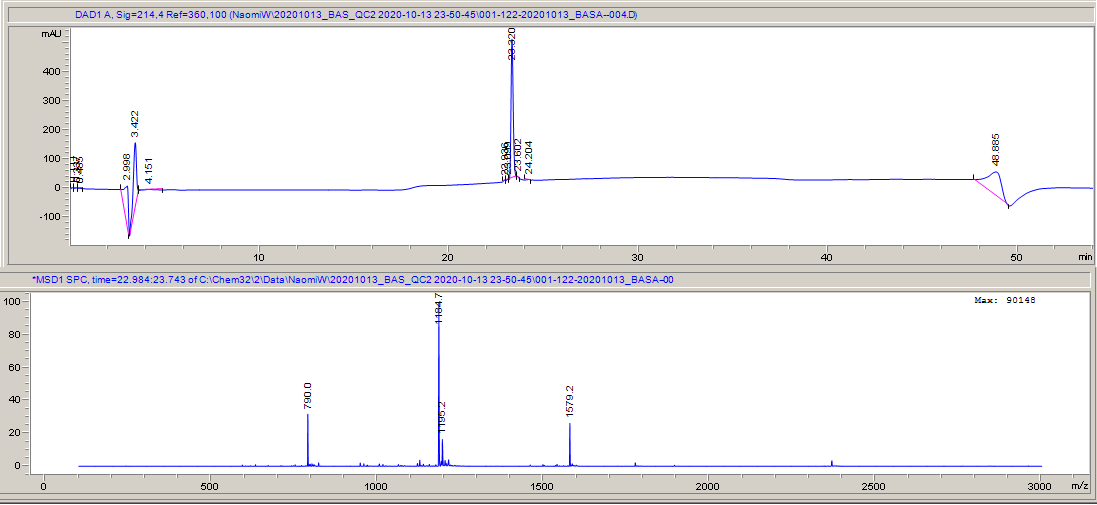

**E.** BTK synthetic substrate F (DEFHMIYEAGVGGK_BIOTIN_GG)

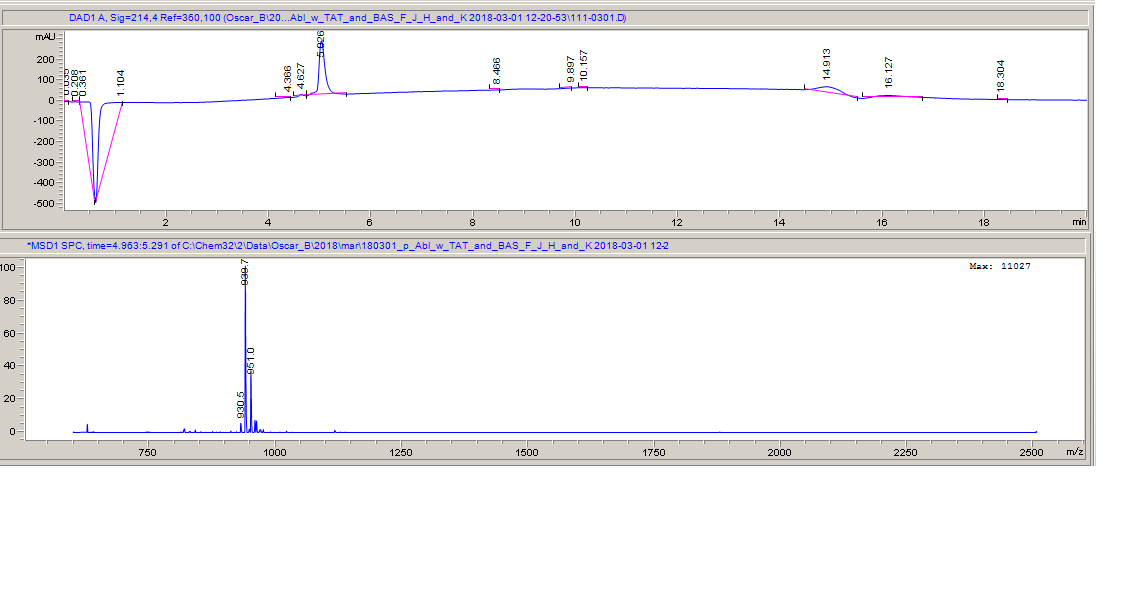
**F.** BTK synthetic substrate H (GGDKQLYATVKGGK_BIOTIN_GG)

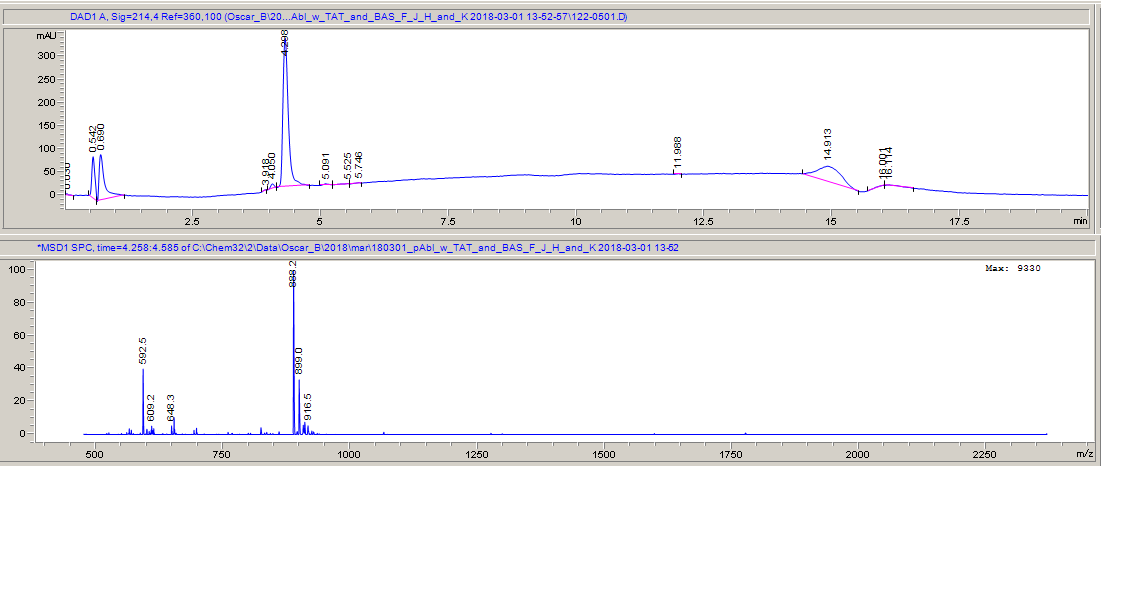

**G.** BTK synthetic substrate K (GGEDDFQMYEQVKGGK_BIOTIN_GG)

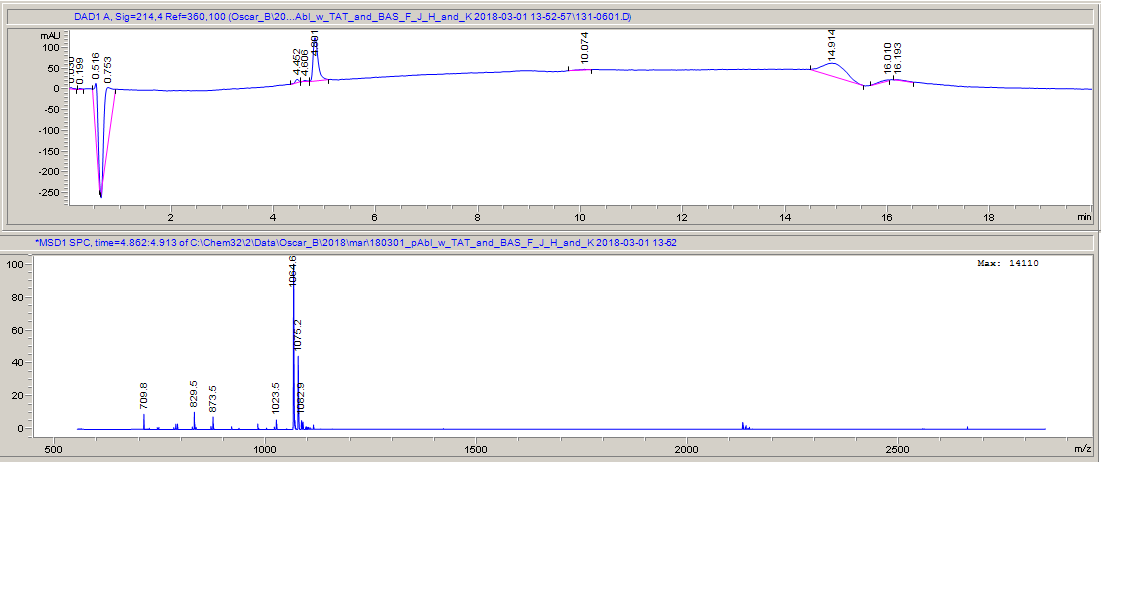

**H.** BTK synthetic substrate T.1 (GGDPDEQIYEGKKEEGGK_BIOTIN_GG)

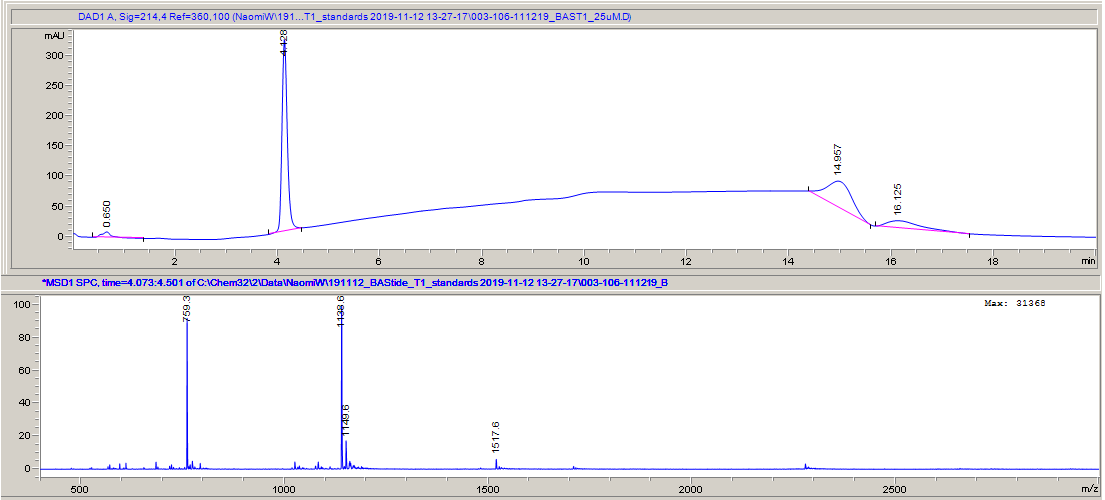

**I.** BTK synthetic substrate phospho-T.1 (GGDPDEQIpYEGKKEEGGK_BIOTIN_GG)

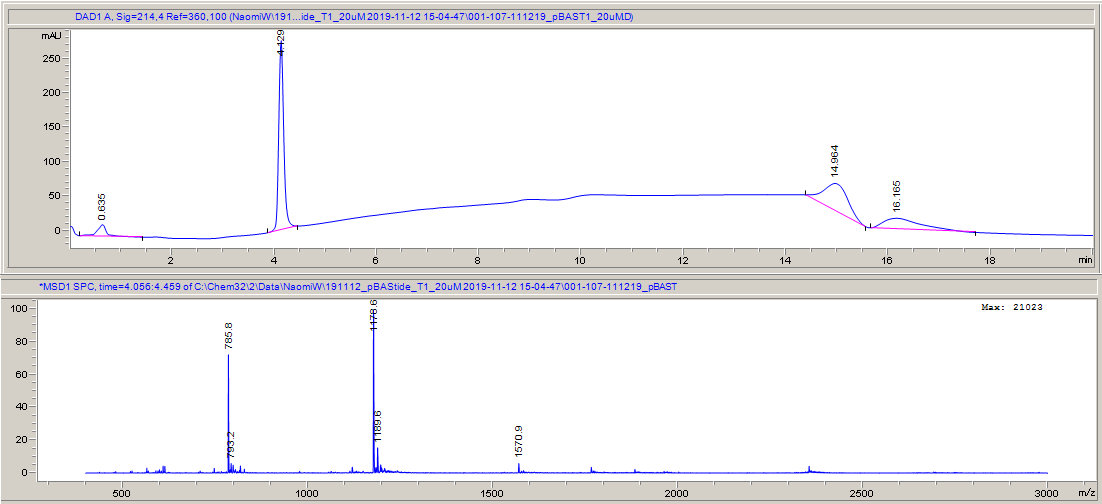

**J.** BTK synthetic substrate T.2 (GGPDDQIYEGKKEGGK_BIOTIN_GG)

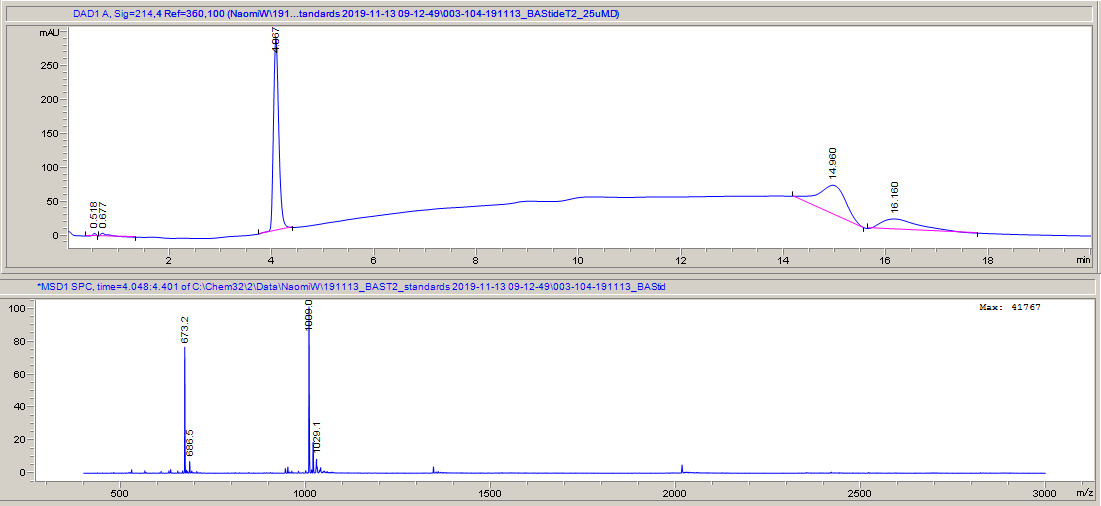

**K.** BTK synthetic substrate phospho-T.2 (GGPDDQIpYEGKKEGGK_BIOTIN_GG)

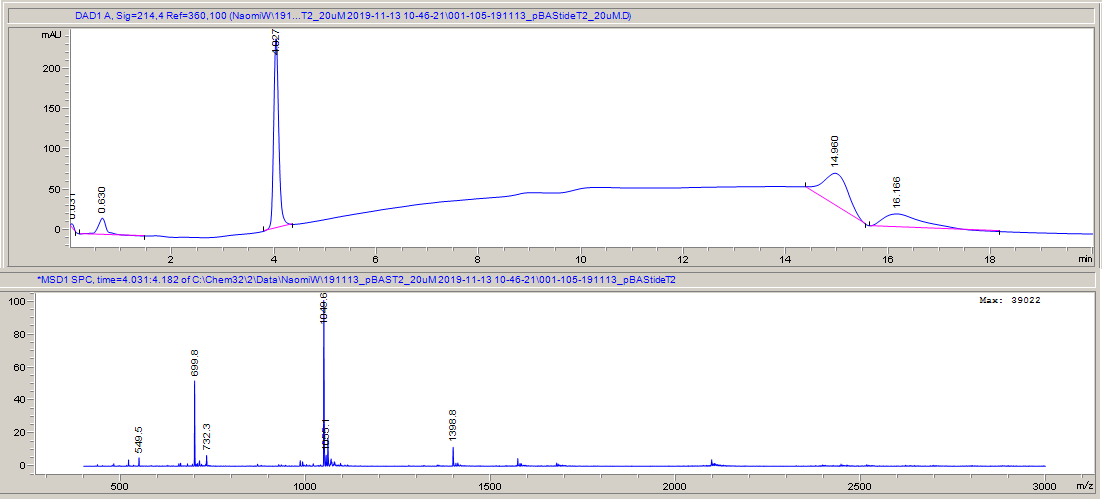

**L.** BTK synthetic substrate T.3 (GGDPDEQIYEEKKEEGGK_BIOTIN_GG)

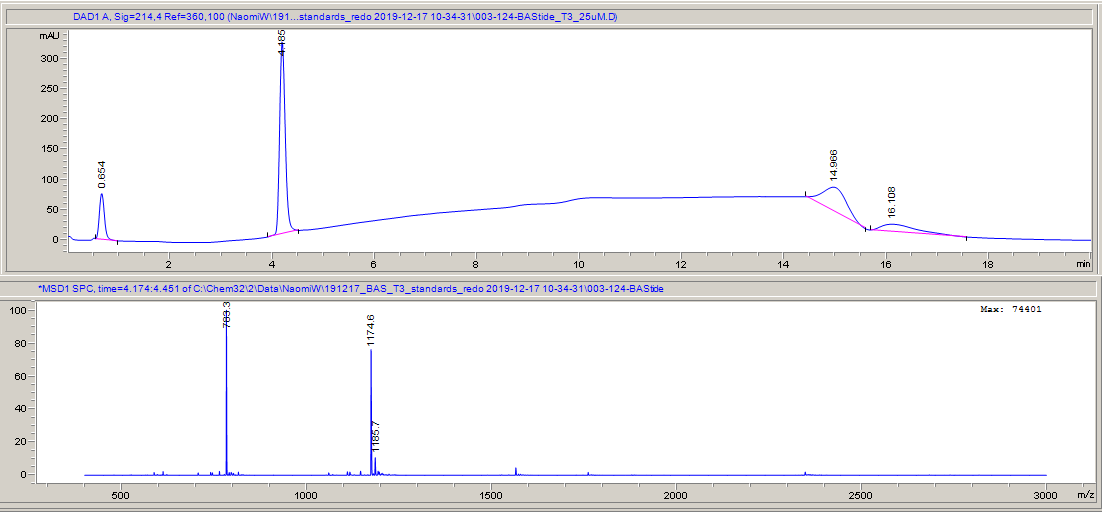

**M.** BTK synthetic substrate phosho-T.3 (GGDPDEQIpYEEKKEEGGK_BIOTIN_GG)

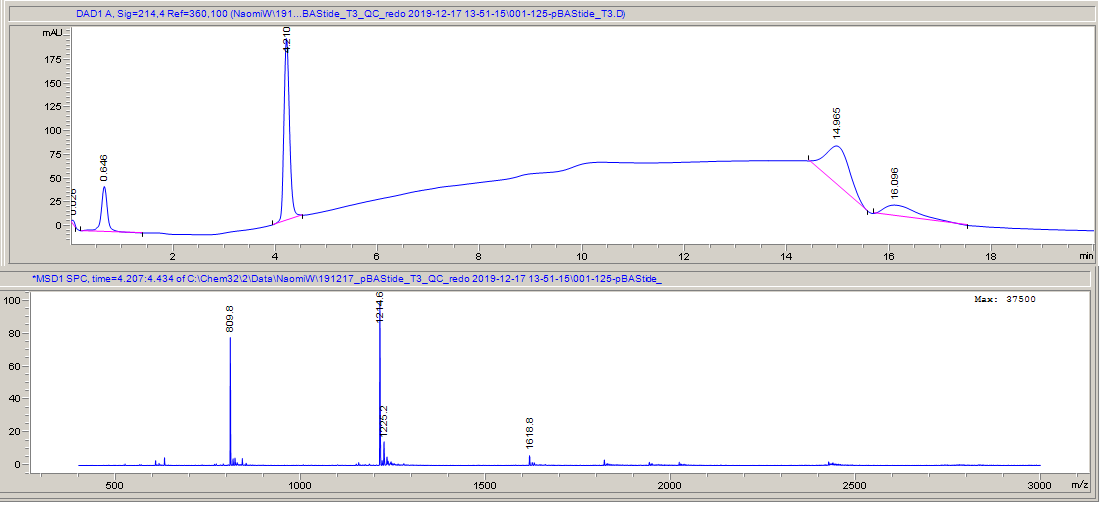

**N.** BTK synthetic substrate T.4 (GGEDPDEQIYEEKKEEEGGK_BIOTIN_GG)

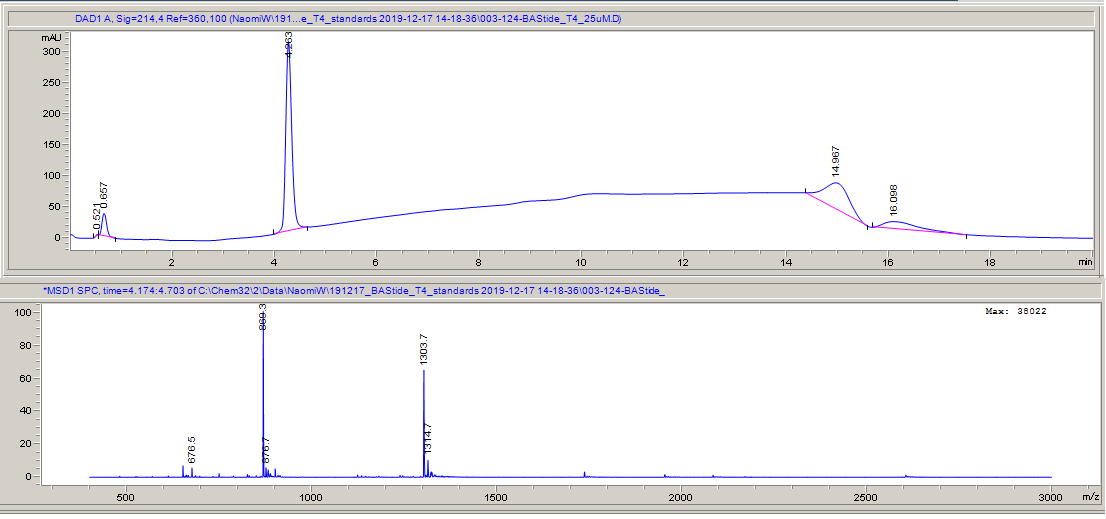

**O.** BTK synthetic substrate phospho-T.4 (GGEDPDEQIpYEEKKEEEGGK_BIOTIN_GG)

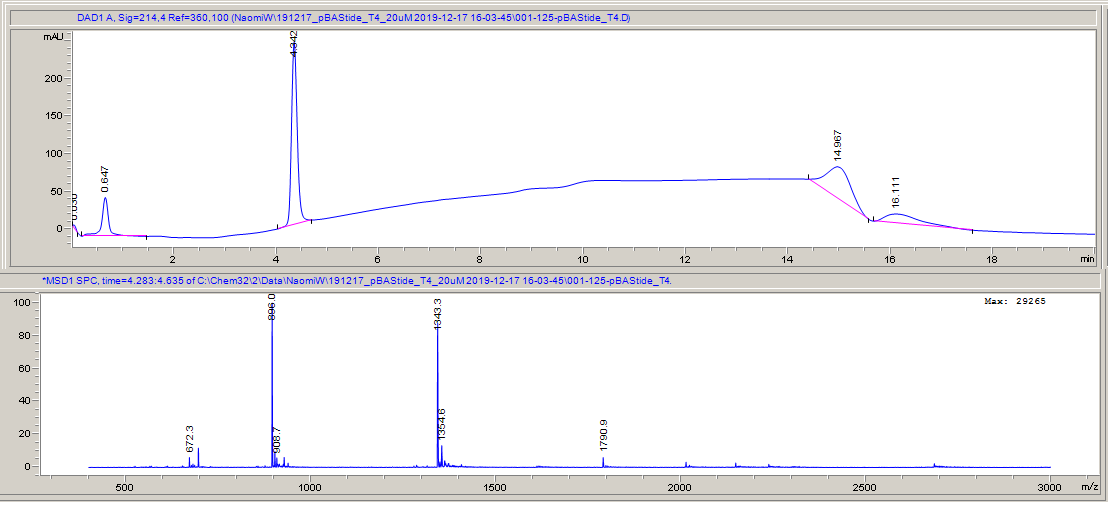
