## Supplementary figures and images for "Novel Bruton’s Tyrosine Kinase (BTK) substrates for time-resolved luminescence assays"

### roc.jpg

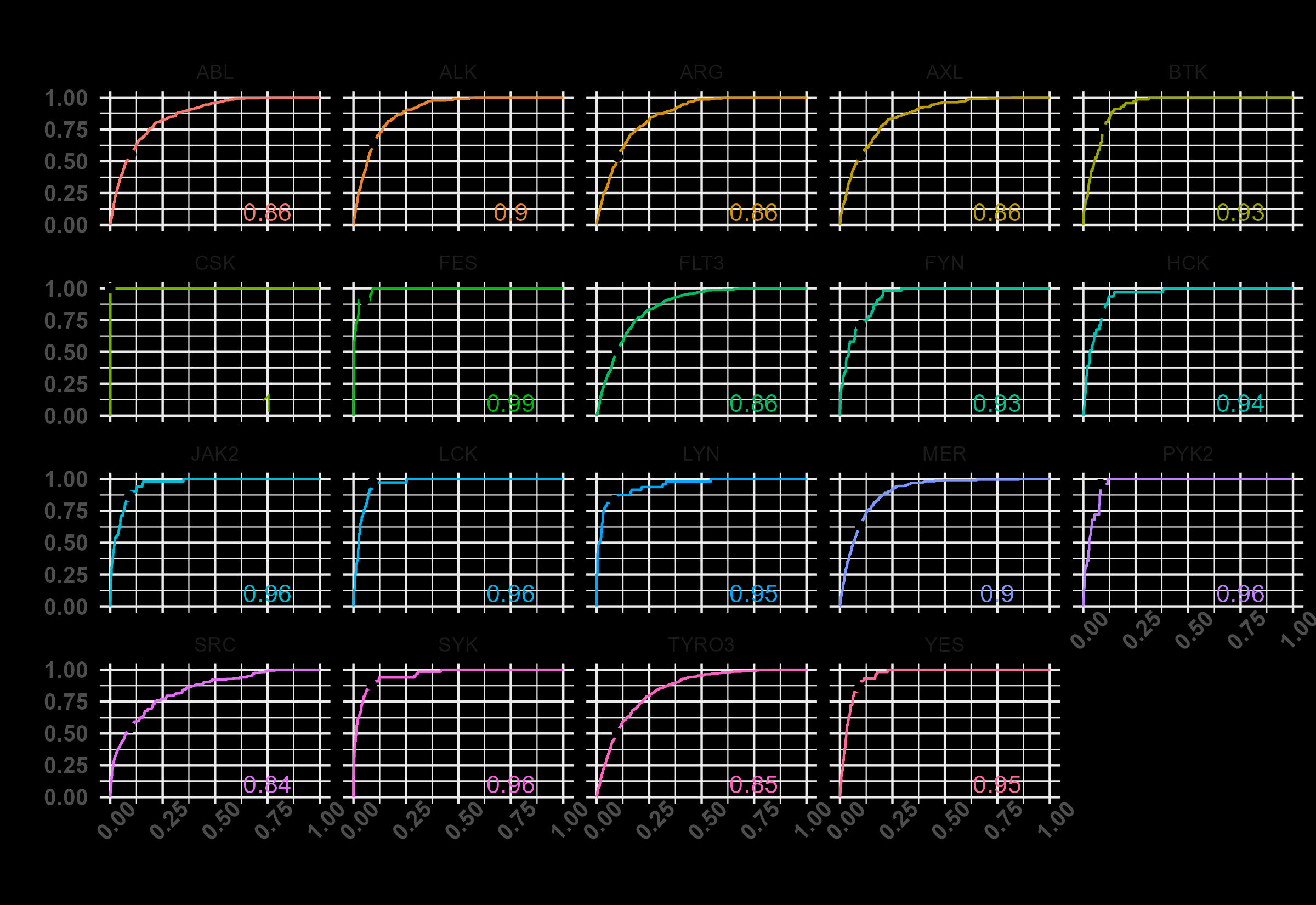

### screener_cutpoints.jpg

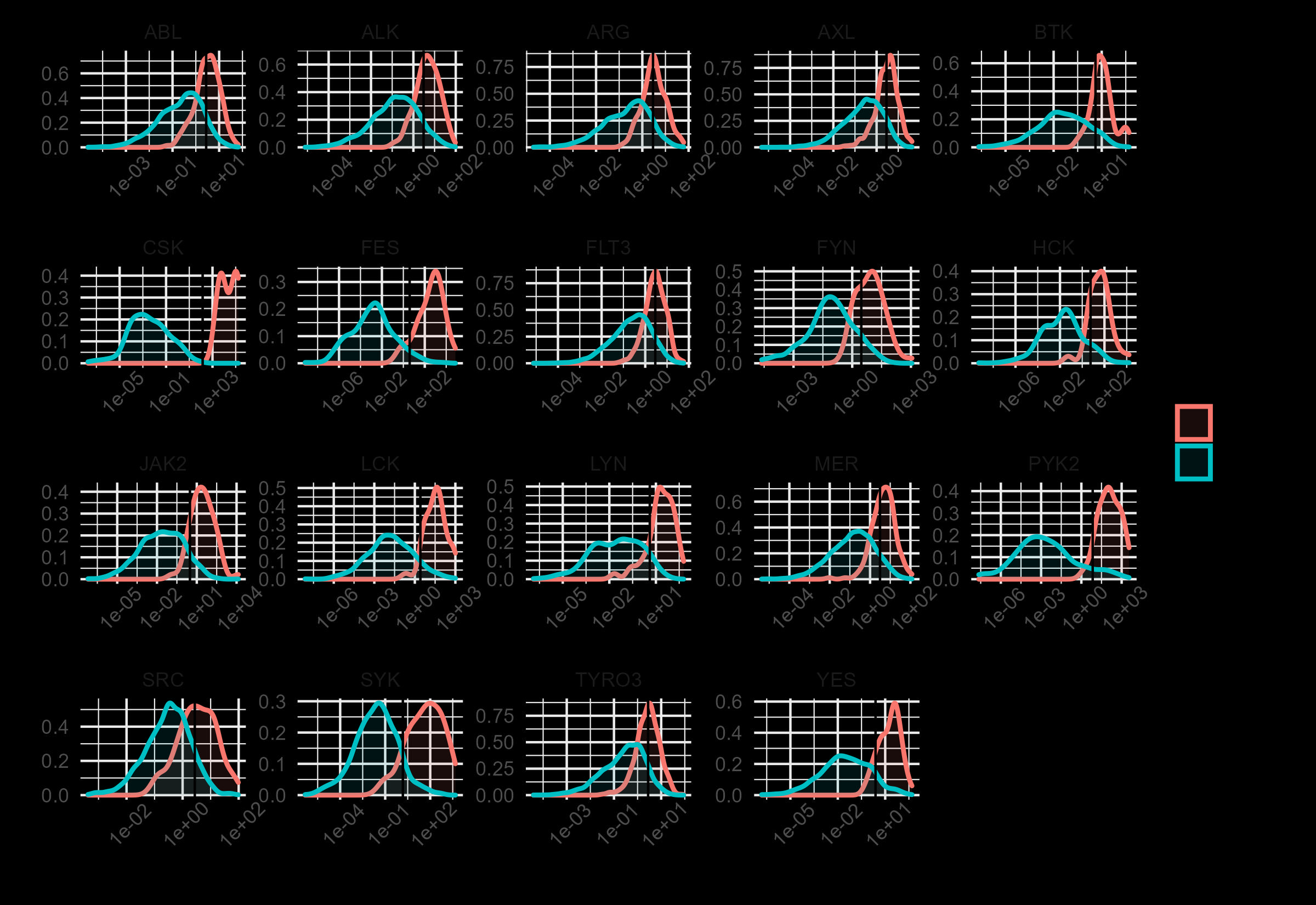
